# Behavioural type drives discovery and exploitation of anthropogenic resources

**DOI:** 10.64898/2026.07.31.742110

**Authors:** Sonja Wild, Isaac K. Uyehara, Lucy M. Todd, Tia A. Ravara, Andrew Sih, Jennifer E. Smith

## Abstract

Human-influenced environments pose challenges but also provide wildlife with anthropogenic resources. Individuals vary widely in their ability to exploit such resources, often as a function of behavioural type. However, we lack a clear understanding of how variation in behavioural traits influences stages of resource exploitation required to use anthropogenic resources. Using fully automated foraging puzzles, we examined how boldness and sociability influenced three aspects of resource exploitation – discovery, problem-solving, and overall performance – in two wild populations of California ground squirrels (*Otospermophilus beecheyi*). Bolder individuals discovered the resource earlier, solved the task faster and achieved higher performance, indicating that boldness promotes efficient exploitation of anthropogenic resources across multiple stages in the process. Greater sociability and more opportunities to observe conspecifics solving the task led to faster problem-solving, consistent with evidence for observational learning. Squirrels in the recreational-use population – with regular exposure to humans and anthropogenic food – were faster to discover the resource than those in a trail-use population – where human exposure was transient and no anthropogenic food available. Problem-solving latency and performance were consistent between populations. Our findings highlight how individual variation in behavioural traits drives performance in novel ecological contexts, providing a mechanistic understanding of behavioural plasticity in human-influenced environments.

## Background

Many animals in the modern world live alongside humans. Although human-influenced environments pose numerous challenges, anthropogenic resources such as shelter or food also provide wildlife with ecological opportunities [1,2]. Locating and exploiting these resources is key to co-existence alongside humans [3,4]. Here, we test a framework that examines multiple factors influencing different behavioural aspects related to exploitation of anthropogenic resources in nature, from initial discovery to subsequent problem-solving to final task performance.

Importantly, there is substantial among-individual variation in animals’ ability to locate, access and utilize anthropogenic resources, much of which has been attributed to consistent individual differences in behaviours across time (personalities) [5–7]. Because these behavioural types can influence how individuals perceive, sample, and respond to their environment [8,9], they are also expected to shape cognitive processes that underlie learning and problem-solving processes, and thus influence the ability to exploit novel resources in human-influenced environments [10,11].

Urban environments in particular often favour proactive (‘faster’) behavioural types that are less neophobic (fearful of novelty [12,13] but are instead characterized by increased *boldness* (higher propensity to take risks), and *exploratory tendencies.* Such patterns have been documented across a wide range of taxa, including coyotes [14,15], ground beetles [16] and song sparrows [17]. Proactive behavioural types are frequently associated with accelerated discovery of and increased engagement with novel resources [8,18]. However, discovery and interacting with a resource alone may be insufficient for its exploitation if anthropogenic resources also require animals to further solve the problem of accessing its concealed contents, such as by opening the lid of a trash bin [19] or piercing milk bottle lids [20].

Consequently, urban environments may not only favour proactive behavioural types, but also select for enhanced problem-solving abilities [10,21–23]. One potential explanation is that behavioural traits such as boldness may be functionally linked to cognitive processes through shared effects on information gathering, persistence, learning and decision-making (‘cognitive style’: [8,9]). However, empirical evidence linking behavioural types and cognitive performance is mixed [24–28].

Beyond boldness, *sociability* – i.e., an individual’s tendency to associate with other individuals [29] – may also play a critical role in resource discovery and exploitation in human-influenced environments. Increased sociability can facilitate faster discovery of anthropogenic resources through improved access to social information [30–32] and may promote interactions with unfamiliar objects via local or stimulus enhancement when conspecifics are present [33]. In addition, opportunities for observational learning can substantially increase the efficiency with which individuals solve novel problems, accelerating the rate at which they learn to access and exploit anthropogenic resources [19,34]. Conversely, sociability may negatively affect resource discovery (e.g. [35]) and exploitation when presence of conspecifics leads to competition, interference or social inertia, preventing access to the resource and/or problem-solving [32,36].

Despite growing recognition that behavioural types play a key role in shaping how animals interact with anthropogenic resources, we still lack a clear understanding of how behavioural traits influence different aspects of resource use. Individuals may vary not only in their propensity to explore environments and approach anthropogenic resources, but also in their ability to solve problems once resources are encountered [37]. Although all aspects are critical for resource exploitation, few studies have simultaneously measured how multiple behavioural axes contribute to the discovery, problem-solving and task performance of wild animals encountering an anthropogenic resource. Understanding how variation in behavioural types shapes all three phases of resource exploitation is therefore crucial for identifying the mechanisms that allow animals to thrive in rapidly changing, human-altered landscapes.

Here, we investigate how variation in two behavioural traits – boldness and sociability – affects the discovery and exploitation of anthropogenic resources in a native mammal that has successfully persisted in human-influenced environments, the California ground squirrel (*Otospermophilus beecheyi*). This facultatively social, opportunistic and behaviourally flexible species regularly exploits human-provided resources [38–42], and exhibits consistent individual differences in behaviour [43,44]. These characteristics make this species an ideal system for examining the mechanisms by which variation in behavioural type influences the discovery and exploitation of anthropogenic resources.

We deployed automated foraging puzzles with a horizontal lever in two populations of squirrels that were ecologically similar but exposed to different levels and forms of human and domestic dog activity [39,41,45]. We used these devices to quantify how individual variation in boldness and sociability affects three aspects of resource exploitation: (i) latency to discovery, (ii) latency to problem-solving, and (iii) task performance (problem-solving rates). We predicted that bold individuals would discover and learn to solve puzzles faster than less bold squirrels [8]. In addition, we predicted that increased sociability might lead to faster resource discovery and problem-solving due to greater availability of social information and possibly a local enhancement effect but negatively affect overall performance due to increased potential for competition and interference by conspecifics. Finally, we predicted that squirrels from the recreational-use population – characterized by frequent exposure to anthropogenic food through picnickers – would be faster to discover and learn to solve the puzzle box compared to squirrels from the trail-use population, where humans were primarily encountered as transient hikers and anthropogenic food opportunities were limited.

## Methods

### Field methods

#### Study sites and live trapping

This research was conducted from May to August of 2024 as part of our long-term study on the behavioural ecology of California ground squirrels at Briones Regional Park, California, United States (37.956798, -122.124437, WGS 84). Experimental subjects belonged to one of two study populations that were exposed to different levels and forms of human activity [39]. The ‘recreational-use’ population was situated in an area with picnic tables and an outhouse, and was characterized by frequent exposure to humans, dogs and anthropogenic food sources through picnickers [41]. In contrast, the ‘trail-use’ population was subjected to less frequent and more transient exposure to hikers, bikers and dog-walkers, with no access to anthropogenic food sources [39]. The two populations were exposed to the same weather conditions, quantified as seasonal patterns in rainfall, temperature and relative humidity [46] (Figure 1b). Individuals from the recreational-use population were generally less fearful of humans, have higher glucocorticoid levels, and were in poorer body condition than those from the trail use population [39,45]. Movement between the two populations was minimal, with roughly 2 % of individuals permanently dispersing between sites [45]. On a weekly basis (typically three consecutive days), we live-trapped and marked individuals using 58 Tomahawk live traps (Hazelhurst, Wisconsin, USA) baited with sunflower seeds and peanut butter. We alternated trapping effort between the two populations. Following established methods at this site, we recorded fear responses (i.e., alarm calls, chatters, and/or struggling) of each trapped squirrel to an approaching member of the field crew [39,47]. Each squirrel was subsequently transferred to a cloth bag to minimize stress while handling [48]. Upon its first capture, we marked each individual with a uniquely numbered Monel ear tag (National Band and Tag., Co, Newport, Kentucky, USA) in one pinna, a subcutaneous PIT-tag (Biomark Inc., Boise, Idaho), and a unique fur dye mark using Nyanzol cattle dye. All individuals were sexed (male, female), aged (adult, juvenile) based on body size and reproductive status, and weighed before being released at the site of capture.

**Figure 1:**
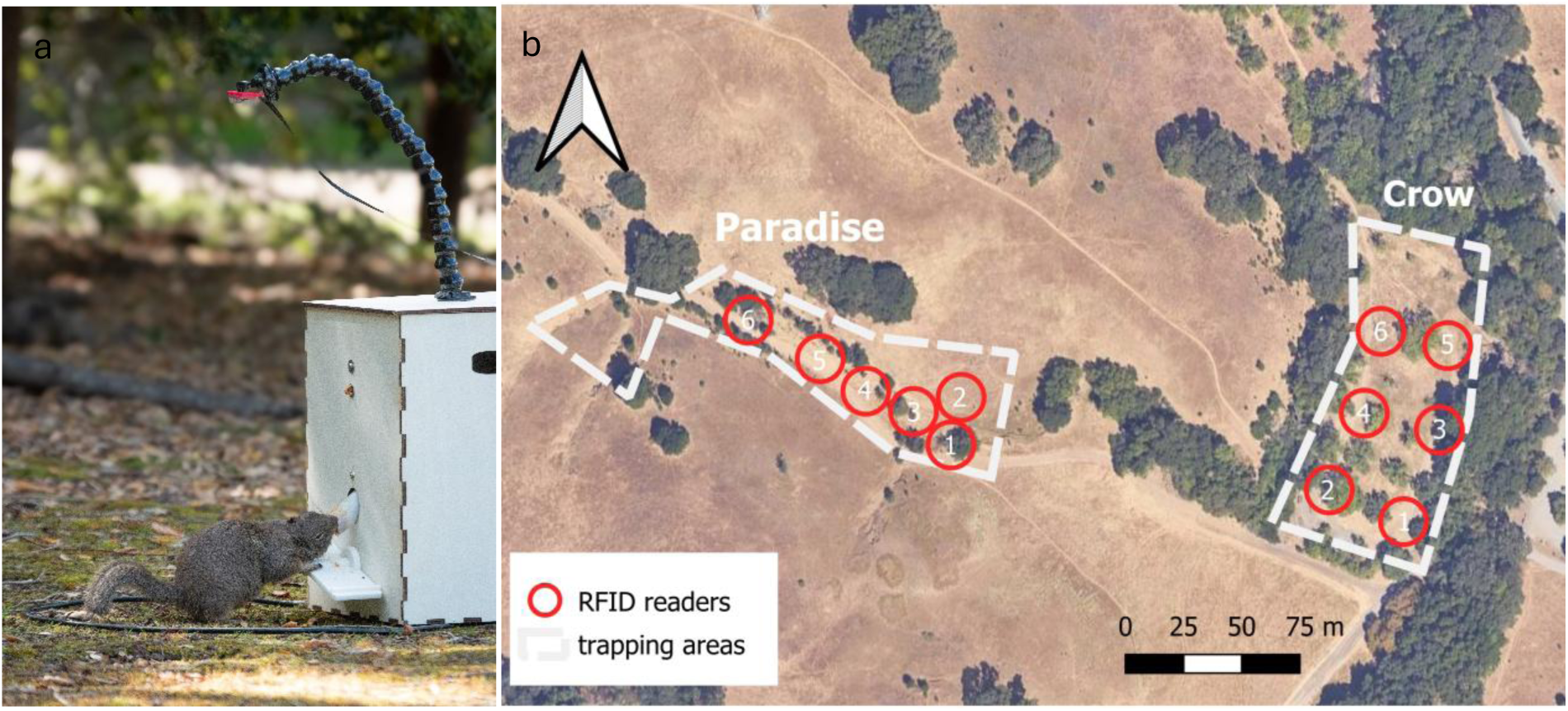
Puzzle box experiments in California ground squirrels. **a: Fully automated experimental puzzle box.** To obtain a food reward, squirrels had to step on a horizontal lever. An RFID loop antenna in front of the box registered squirrel identity via its unique PIT tag. **b: Study area with twelve experimental locations across two populations.** The recreational-use population to the east was subject to moderate exposure to anthropogenic activity from domestic dogs, picnickers and other recreational activities, whereas the trail-use population to the west was subject to occasional human foot traffic, bicyclists, and dogs. Image credit: Satellite basemap from Esri World Imagery.

#### Puzzle box experiments

To investigate how variation in behavioural types (i.e., boldness and sociability) predicted exploitation of anthropogenic resources, we deployed custom-built and fully automated puzzle boxes (see SI for detailed description of hard- and software; Figure 1a; Figure 2b-c) on days when traps were not set. To obtain a small number of sunflower seeds and kibbled peanuts – which are a highly desirable food reward [49] – squirrels had to step on the right or left side of a horizontal lever at the front of the puzzle box and rotate the lever by at least 15°. The puzzle box automatically logged the identity and time each squirrel entered (and departed) the antenna area in front of the box via its PIT-tag using RFID techniques. It also logged the time and solving side (right or left) if a solve occurred.

**Figure 2:**
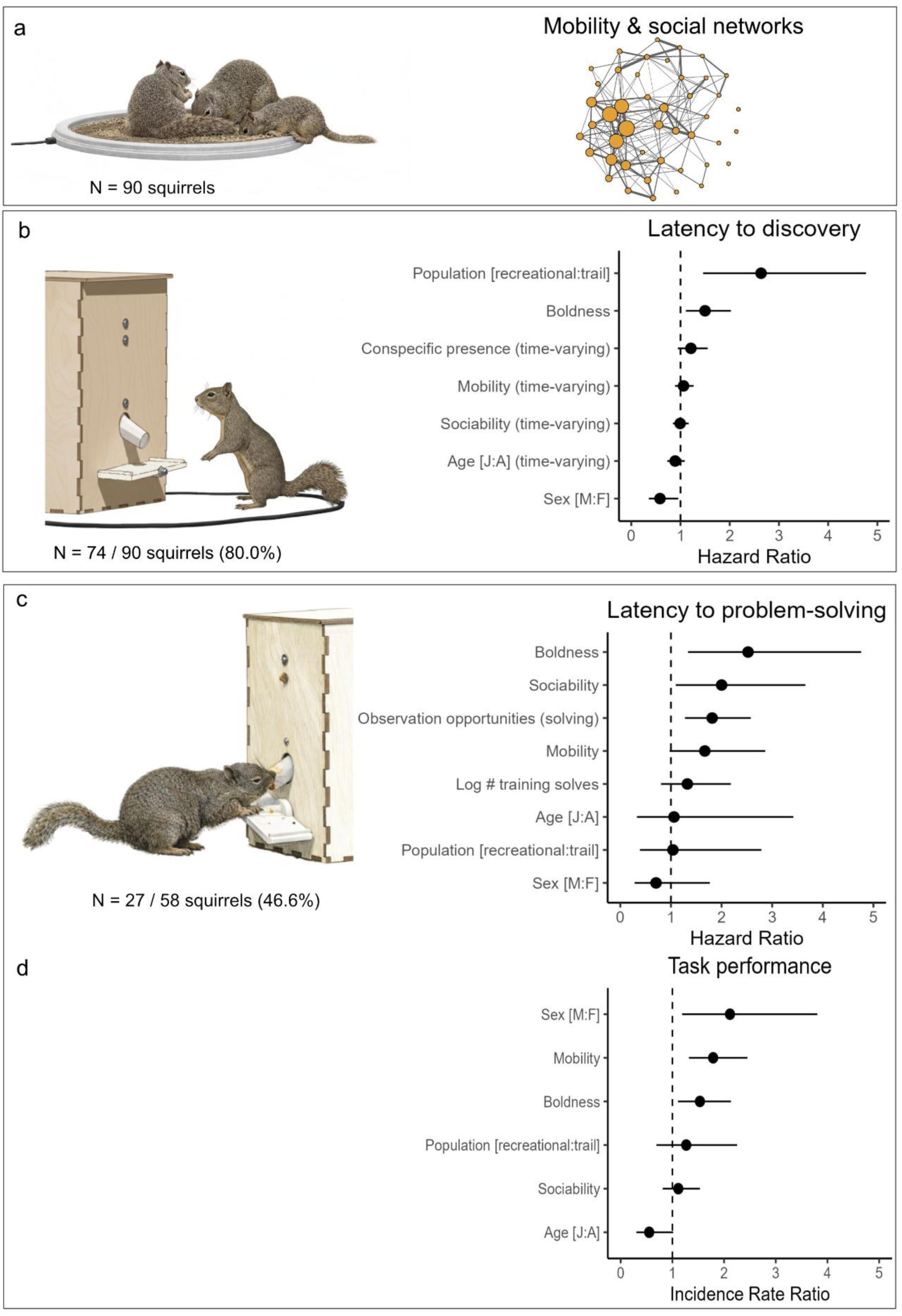
Variation in behavioral type affects exploitation of anthropogenic resources in California ground squirrels. **a: Mapping space use and social networks.** Space use and co-feeding events were measured using grids of baited RFID antennae to quantify individual mobility and eigenvector centrality as a measure of sociability. **b: Latency to discovery of experimental puzzle boxes.** Squirrels from the recreational-use population and bolder individuals were faster to discover the anthropogenic resource while males were slower relative to females. **c: Latency to problem-solving.** Boldness, sociability, mobility and the number of observation opportunities of conspecifics solving were connected to faster problem-solving. **d: Performance.** Males, more mobile and bolder individuals had higher solving rates while juveniles had lower rates relative to adults. Illustrations were generated using AI-tools from original photographs by the authors, which are provided in Figure 1a and in Figure S3, respectively. Error bars represent S5% confidence intervals.

Puzzle box experiments consisted of a training phase (May 8-June 27) and trial phase (July 10-August 6). The training phase was designed to encourage squirrels to approach the unfamiliar apparatus and allow a subset of individuals to learn the association between lever manipulation and food reward to serve as potential demonstrators for conspecifics during the subsequent trial phase. To facilitate interaction with the puzzle box, a small amount of peanut butter was added to both sides of the horizontal leaver, and food rewards were provided following successful lever manipulation.

At each training session, we simultaneously deployed three puzzle boxes spaced approximately 20 m apart at one of six experimental locations within one of the two populations (Figure 1b). Experimental locations were distributed across the study area to ensure that all individuals had access to the puzzle boxes within their home range. Individuals visited the puzzle boxes at between one and six locations within their population (mean = 2.4 locations). Deploying multiple puzzle boxes simultaneously reduced the likelihood that individuals would monopolize access to the apparatus and increased opportunity for nearby squirrels to interact with the resource. Training sessions lasted 2-3 h (mean: 190 minutes; S.E: ± 17 min) and continued at a location until at least 10 successful solves had occurred, irrespective of squirrel identity.

Training effort differed between populations. In the recreational-use population, the criterion of 10 solves was reached after 9h 45 minutes, whereas the trail-use population required 25h and 13 minutes to reach the same criterion. The potential effect of individual differences in exposure to the puzzle box during training on problem-solving latency and performance (solving rate) during the trial phase was accounted for in statistical analyses (see below).

During the trial phase, puzzle boxes were deployed without additional peanut butter. On each experimental day, three puzzle boxes were consecutively deployed at three of six locations to ensure that individuals throughout the study area had opportunities to encounter the resource. The order of locations was semi-randomized to avoid prolonged sun exposure and overheating of electronics. At each location, the three boxes were spaced approximately 20m apart and remained deployed for 1h before being moved to the next location, resulting in 3h of exposure per experimental day. Each of the six locations in both populations was sampled 3-4 times, resulting in a total of 21 experimental hours per population.

#### Quantifying behavioural types and space use

For each squirrel in the two populations, we extracted measures for two behavioural traits that could conceivably influence exploitation of anthropogenic resources, namely (i) boldness; and (ii) sociability (Figure S2).

To compute *boldness* scores, we ran a principal component analysis (PCA) [50] on individuals’ trappability (proportion of trapping days for a given population where a focal individual was trapped; bolder animals are trapped more often), and the proportion of trapping events during which the squirrel expressed one or more fear responses (chatter, call, struggle). These behaviours have previously been used to assess consistent individual differences in these populations [39,43]. The two variables loaded equally and in opposite directions on PC1 (trappability: 0.827; fear responses: -0.827) where PC1 accounted for 68.4% of the variance. Four individuals from the recreational-use population and eight individuals from the trail-use population were detected by the RFID antennae but not trapped in summer 2024. These were assigned the minimum boldness score resulting from the PCA.

To quantify individuals’ sociability, we recorded co-feeding events during a separate RFID-based assay independent of the puzzle box experiments. In this assay, we simultaneously deployed three clusters of RFID readers, each with three baited 0.25m loop antennae (Biomark Inc.) at three out of the six experimental locations (Figure 1b; Figure 2a). The RFID readers continuously logged the presence of PIT-tagged individuals within each antenna’s detection range, recording individual visits with high temporal resolution. PIT-tag detections were sequential and multiple individuals could be recorded by the same antenna in rapid succession.

After 45 minutes, all antennae were rebaited with sunflower seed and left for another 45 minutes, before being moved to the second set of three experimental locations, where the same procedure was repeated (45 min feeding, rebaiting, 45 min feeding). This design ensured complete spatial coverage of each population within a single experimental day. The procedure was then repeated in the second population on the following day, and data were collected approximately once per week per population across five sampling days.

To obtain measures of *sociability*, we first computed co-feeding events from the fine-scale spatiotemporal stream of RFID detections using Gaussian mixture models implemented in the R package ‘asnipe’ [51]. These methods identify temporally overlapping visits to the same antenna as social co-occurrence events. We then built dyadic association networks using the simple ratio index which ranges from 0 (never associated) to 1 (always associated) [52,53]. From these networks, we extracted individual eigenvector centrality as our measure of sociability, which is the sum of centralities of an individual’s neighbours and captures the potential of individuals to serve as social hubs or for the spread of social information [54].

Finally, since space use can influence sociality [55] and the probability of discovery and encounter frequency of resources, we computed a measure of *mobility* within the study area as the median number of distinct RFID antennae visited per experimental day (out of total 18 possible).

### Statistical analyses

We fit three separate models to investigate factors predicting (i) latency to discovery; (ii) latency to problem-solving and (iii) task performance. Latencies to discovery and problem-solving were analysed using Cox proportional hazard models using R package ‘survival’ [56]. Survival models are used to model time-to-event data and estimate a hazard rate for each covariate as a ratio relative to the reference level [57]. Hazard ratios greater than 1 indicate a higher hazard (shorter latency to the event), whereas values below 1 indicate a lower hazard (longer latency). Individuals that did not discover or solve the task by the end of the experiment in their respective population were right censored, which accounts for differing lengths of the training period between the two populations.

For each model, we assessed the proportional hazards assumption, which requires that covariate effects remain constant over time [58]. When predictors violated this assumption, they were modelled as time-varying effects by including an interaction with log(time), following standard practice for accommodating non-proportional hazards in Cox models [59].

#### Latency to discovery

For modelling the latency to discovery, we included squirrels that had either been registered on RFID readers or puzzle boxes at any point during either the training or trial phase. Discovery latency was defined as the time elapsed between first deployment of puzzle boxes within a population and an individual’s first arrival at any puzzle box. Two individuals that discovered puzzle boxes in both populations were retained only in the population where they made their first discovery. As predictor variables, we included measures of (i) boldness; (ii) eigenvector centrality (iii) mobility; and additionally controlled for (iv) population; (v) age category (juvenile vs adult); (vi) sex (male vs female); and (vii) presence of a conspecific within the antenna area at time of discovery (binary variable). If the latter is a positive effect it accounts for effects of stimulus or local enhancement; i.e., when the presence of a conspecific increases the probability of a naïve squirrel approaching or interacting with an object, and is a form of social information use [33]. Alternatively, if it is a negative effect, the presence of conspecifics likely reflects competition or interference. A conspecific was considered as present at the box, if it was registered by the antenna within the 10s preceding the arrival of the focal individual. Given typical movement patterns, detections within this window meant that the individual was likely still within ∼5m of the box – a biologically meaningful distance for gathering and reacting to visual information in this study species [60,61].

#### Latency to problem-solving

For modelling the latency to problem-solving, we included all individuals that had discovered the puzzle box, including individuals first detected during the trial phase. However, analyses of problem-solving latency were restricted to the trial period because peanut butter applied directly to the lever during training increased the likelihood of incidental solves while squirrels explored or attempted to access the peanut butter. Although accidental lever depressions could still occur during the trial phase, repeated successful solves were more likely to reflect consistent performance rather than chance interactions with the apparatus. We therefore classified individuals as ‘knowledgeable’ once they had achieved at least three successful solves during the trial phase and defined problem-solving latency as the time elapsed between the start of the trial phase and an individual’s third successful solve. Naïve squirrels that produced fewer than three solves were included as right censored. We again included predictors for (i) boldness; (ii) sociability; (iii) mobility; (iv) population; (v) age category; (vi) sex; and (vii) the log number of solves produced during the training phase to control for opportunities for trial-and-error learning prior to the trial phase; and (viii) the total number of observation opportunities of conspecifics solving. The latter served to assess a potential effect of observational (social) learning. We assumed that a focal individual had opportunity to observe another conspecific solve the puzzle if the focal was registered by the antenna of the same box within 10s of a solve occurring.

#### Task performance

To model performance, for knowledgeable squirrels that had solved puzzles at least three times, we extracted the total number of solves they produced. This was included as an outcome variable in a generalized linear model with a negative binomial error distribution to account for overdispersion in solve counts [62]. The number of solves was modelled as a function of (i) boldness; (ii) sociability; (iii) mobility; (iv) population; (v) age category; and (vi) sex; with the log-transformed number of experimental days on which squirrels were detected at the puzzle boxes after learning to solve as an offset to control for individual differences in exposure to the task such that performance was measured as solving rates per experimental day.

We assessed variance inflation factors and correlations among behavioural predictors prior to analyses, and tested for population-level differences in boldness, sociability and mobility using Mann-Whitney U tests.

## Results

During the field season, we trapped and marked a total of 142 individual squirrels (100 in the recreational-use population, 42 in the trail-use population). Mobility and sociability were moderately correlated (r = 0.73), whereas boldness showed only weak correlations with the other traits (|r| ≤ 0.21). Variance inflation factors were low for all predictors (all VIFs < 2.3; Table S1), indicating that collinearity was not substantial. We therefore retained mobility and sociability as separate variables because they capture biologically distinct aspects of behaviour. Squirrels from the recreational-use population scored higher in boldness than those from the trail-use population (Mann Whitney U: U=1276; p=0.007). However, we did not detect population-level differences in mobility (Mann Whitney U: U=1125; p=0.145) or sociability (Mann Whitney U: U=1060; p=0.362; Figure S2).

### Latency to discovery

Overall, 90 squirrels were present at either study area during the training or trial phase, of which were 36 adult and 16 juvenile females, 25 adult and 13 juvenile males. Of those, 72 (80.0%) discovered the puzzle boxes – 32 adult and 13 juvenile females, 19 adult and 8 juvenile males. Squirrels from the recreational-use population were faster to discover the novel resource (hazard ratio (HR) = 2.64 [1.46-4.77], p = 0.001) compared to squirrels from the trail-use population (Figure 2b). Even after controlling for site-level effects, boldness still significantly decreased the latency to discovery (HR = 1.50, 95% CI = [1.11, 2.02], p = 0.008), and males had higher latency than females (HR = 0.58, 95% CI = [0.36, 0.95], p = 0.032). No significant effects were detected for sociability, mobility, age, or conspecific presence (all p > 0.11; Table S2). The overall model fit was good (likelihood ratio test: χ² = 36.07, df = 7, p < 0.001), with a concordance of 0.707.

### Latency to problem-solving

In total, 27 out of 58 squirrels present at either site during the trial phase (46.6%) learned to solve the box (i.e., produced a minimum of 3 solves). Of the 27, 17 were female (9 adults and 8 juveniles) and 10 male (9 adults and 1 juvenile). Boldness significantly decreased the latency to solve (HR = 2.52, 95% CI = [1.34, 4.76], p = 0.004), as did sociability (HR = 2.00, 95% CI = [1.10, 3.66], p = 0.024), and the number of observation opportunities of conspecifics solving (HR = 1.81, 95% CI = [1.28, 2.58], p > 0.001; Figure 2c). More mobile individuals were marginally faster at learning to solve (HR = 1.67, 95% CI [0.97-2.86], p = 0.062), though the confidence interval slightly overlapped 1. Population, age, sex, and the number of ‘solves’ during the training phase (see Methods) did not significantly influence latency to solve (all p > 0.27; Table S3). Overall model fit was good (likelihood ratio test: χ² = 45.88, df = 8, p < 0.001), with high predictive accuracy (concordance = 0.84).

### Task performance

Across the 27 squirrels who learned to solve the box, boldness significantly increased performance (i.e. daily solving rates) (incidence ratio rate (IRR) = 1.53, 95% CI = [1.11, 2.13], p = 0.004), as did mobility (IRR = 1.79, 95% CI = [1.32-2.45], p < 0.001; Figure 2d). Juveniles had lower solving rates compared to adults (IRR = 0.55, 95% CI [0.30-1.01], p = 0.036), while males had higher rates than females (IRR = 2.11, 95% CI [1.19-3.81], p = 0.009). In contrast, neither sociability nor population predicted performance (all p > 0.41; Table S4).

## Discussion

Using fully automated foraging puzzles, we assessed how variation in behavioural type predicted resource exploitation by members of two populations of California ground squirrels. Going beyond previous studies, we investigated how individuals’ boldness and sociability related to three different aspects of anthropogenic resource exploitation – latency to discovery of the task, latency to problem-solving, and overall performance.

Increased boldness was associated with better success across all three phases of exploiting the anthropogenic resource. Bolder individuals were faster to discover it, faster at learning to access it, and had higher performance (i.e., daily solving rates) than less bold individuals. Higher mobility was associated with faster problem-solving and better performance. These results suggest that ‘fast’ personality types – characterized by increased boldness and movements within sites – were overall fastest to discover and learn to solve puzzles. These findings are consistent with the notion that reduced neophobia and willingness to approach and engage with novel anthropogenic objects is a key component facilitating different aspects of resource exploitation [8,18,63]. Besides leading to faster discovery and engagement, bolder individuals also achieved higher solving rates and were thus more successful at exploiting the resource. This aligns with a large-scale meta-analysis demonstrating an overall link between problem-solving ability and boldness – when boldness is measured as reactivity to (simulated) predation [24].

In our study, boldness may have facilitated exploitation of anthropogenic resources through a generalized tolerance toward humans and human-made objects. Because our measure of boldness integrated both trappability – where traps represent human-made objects associated with food – and reduced fear responses to approaching humans, bolder individuals may have been more habituated to anthropogenic stimuli overall [64]. Such habituation can result in a generalized reduction in fear, where reduced fear toward one class of stimuli (e.g. humans) extends to other, perceptually or contextually similar stimuli, including novel human-made objects (traps, puzzle boxes) [65,66]. As a result, bolder individuals may have been more willing to approach and interact with the foraging task, facilitating both faster discovery and more successful exploitation. Fear generalization as a possible mechanism is further supported by the fact that squirrels from the recreational-use population – scoring overall higher in boldness (Figure S2a) and being more frequently exposed to human activities and anthropogenic food through picnickers [39] – were significantly faster to discover the resource.

Latency to problem-solving was significantly predicted by squirrels’ sociability and opportunities to observe conspecifics solving the puzzles. More sociable individuals with greater exposure to conspecifics solving subsequently solved puzzles faster. While we did not explicitly investigate direct instances of one animal solving immediately after seeing a conspecific solve, our findings are consistent with observational learning [67]. Notably, the effect of sociability remained significant even after accounting for observation opportunities. This suggests that more socially central individuals may not only have had more chances to learn from others but may also have been more attentive to observed behaviours [68,69]. Our findings are consistent with data showing that access to social information facilitates the rapid spread of exploitative behaviours (e.g. bin-raiding in cockatoos: [19]; longline depredation in sperm whales [70]; crop-raiding in African elephants [71]), especially for socially central individuals [72]. In contrast, conspecific presence alone did not influence latency to discovery of the puzzle boxes, suggesting absence of a local or stimulus enhancement effect [33]. Contrary to our expectation, sociability did not influence performance, suggesting that more sociable individuals did not necessarily experience more competition or interference, which could plausibly negatively affect solving rates.

Males were slower to discover the puzzle boxes compared to females, but there was no difference in latency to problem-solving between the sexes. State-dependent sources of motivation such as hunger can greatly increase individuals’ propensity to engage with foraging tasks and lead to higher innovation rates [73–75]. Thus, faster discovery of the puzzle boxes in females relative to males may partly reflect elevated nutritional demands associated with reproduction in late spring [38]. Meanwhile, males had overall higher performance, which could potentially be explained by differential access to defensible resources [76].

Age category did not influence latency to discovery or problem-solving, which stands in contrast to several studies demonstrating that juveniles tend to be less neophobic and more exploratory compared to adults, which should lead to faster discovery of novel resources (reviewed in [77]). Juveniles also had overall lower performance, which could be indicative of developmental constraints on cognitive abilities [28], result from physical limitations of small juveniles when pushing the lever, and/or be explained by reduced access to the puzzle boxes compared to more competitive adults.

Squirrels from the recreational-use population were faster to discover the resource, even after controlling for site-level differences in boldness. Accumulating evidence suggests that different forms of human activity can differentially shape animals’ behavioural responses [78–80]. In our system, squirrels in the recreational-use population were regularly exposed to predictable, non-threatening activities like stationary picnicking, while squirrels in the trail-use population experienced more transient and potentially aversive disturbances associated with hikers, cyclists and dogs. Predictable human presence may promote habituation and tolerance towards humans and anthropogenic objects [81], thereby increasing willingness to investigate novel resources. In contrast, more unpredictable or disturbance-associated activities may suppress exploratory behaviour and reduce opportunities for social information use through altered network connections [40,82]. Of course, additional replication of sites would be required to make definitive conclusions about the mechanisms shaping site-level differences in behaviour. Future studies incorporating fine-scale quantification of human disturbance could determine how both the intensity and nature of human activity interact with individual behavioural type to shape risk perception and, in turn, the cognitive processes underlying resource discovery and exploitation [41,80,83–85].

Importantly, population-level differences were only apparent in the earliest stage of resource use – initial discovery – rather than subsequent performance. Once squirrels had discovered the task, neither latency to problem-solving nor solving performance differed between populations. However, because the two populations differed in training duration prior to the trial phase (see Methods), comparisons of problem-solving latency should be interpreted with caution. In addition, analyses of problem-solving were necessarily conditional on discovery, excluding individuals that never engaged with the resource. If exposure to humans or the nature of anthropogenic activity influenced the probability of discovery, population differences in cognitive performance across the full range of behavioural types may therefore have been partially obscured.

The ability to discover, access and exploit anthropogenic resources is central to persistence in human-influenced systems. Our results demonstrate that individual variation in boldness consistently predicts success across different aspects of resource exploitation, providing a mechanistic link between behavioural type and successful co-existence alongside humans. This study highlights how behavioural traits and cognitive processes interact to shape outcomes in novel ecological contexts. Understanding the causes and consequences of such individual variation is critical for predicting population-level responses to expanding human activity in a rapidly changing world, with important implications for wildlife conservation, urban management and the mitigation of human-wildlife conflict [6,10,86].

## Acknowledgements

We thank the numerous students who have contributed to field data collection on the California ground squirrel population at Briones Regional Park, in particular the 2024 field crew Jay Ingbretson, Mackenzie Miner, Ella Oestreicher, Mari Podas, Lupin Teles, Jada Wahl and Haoyue Dong. We thank the staff of the East Bay Regional Park district, USA, especially Doug Bell and Joseph Miller, and the California Department of Fish and Wildlife, USA, for their support and cooperation. We thank the Sih lab for fruitful discussions during preparation of this manuscript.

## Use of Artificial Intelligence (AI) and AI-assisted technologies

AI technology was used for language editing.

## Ethics

All field methods were approved by the Institutional Animal Care and Use Committee of the University of Wisconsin-Eau Claire (#333) and the University of California Davis (#23297), U.S.A., and are consistent with the guidelines of the American Society of Mammalogists for the use of wild mammals in research (87). Research permits were obtained from the California Department of Fish and Wildlife, U.S.A. and the East Bay Regional Park District, U.S.A.

## Data, code and materials

All data and code needed to replicate analyses are available at https://github.com/sonjawild/squirrels_puzzle_box. Supplementary results are available in the Supplementary Information.

## Author contributions

Conceptualization: SW. & JES. Methodology: SW. Software: IKU. Fromal analysis: SW. Investigation: SW, LMT, TAR. Resources: SW, JES, AS. Data curation: SW. Writing-Original draft: SW. Writing – Review C Editing: JES, AS, IUK. Visualization: SW. Supervision, project administration, funding acquisition: SW C JES.

