## Supplementary Information for "Behavioural type drives discovery and exploitation of anthropogenic resources"

### Puzzle box – hardware and software

To investigate how variation in behavioural type influenced resource exploitation in human-influenced environments, we built three fully automated foraging puzzles (‘puzzle box’ hereafter). At the front of a wooden box (30x30x40 cm), at a height of 7 cm off the ground, we installed a 20x7 cm plastic lever, with a protruding plastic funnel 4 cm above the lever through which the food reward was dispensed. When squirrels stepped on the horizontal lever (either left or right), two 3D printed internal cog wheels activated a dispenser wheel with a small hole at the bottom of a container to dispense a mix of approximately 5-8 sunflower seeds and kibbled peanuts through the funnel. The system was fine-tuned so that squirrels only obtained a reward if the lever was pushed at an angle of at least 15 degrees.

The puzzle box was controlled by a Raspberry Pi 4 micro-computer powered by a 30mAh power bank, with commercially sourced electronics attached: Two infrared sensors to register solves, two servo motors for resetting the lever to a neutral position and a Raspberry Pi 3 camera module installed over the front of the puzzle box to record squirrels interacting with the box.

Each of the three puzzle boxes was deployed with a 0.61m cord antenna (BioMark, Inc.) around the front of the box to automatically register arrival and departure of PIT-tagged individuals at the box. All three cord antennae were connected to a single RFID reader (Biomark, Inc.). To enable communication between the RFID reader and the three Raspberry Pis, we established a Bluetooth connection between a fourth (master) Pi and the RFID reader, which sent information to the three box Raspberry Pis about arrival and departures of squirrels in real time via a local Wi-Fi network.

The software was custom-built for the Raspberry Pi (Figure S1). The master Pi communicated with the RFID reader to establish which antennae were reading the presence of a PIT tag and then logged this information in a log file. The other Raspberry Pis requested the RFID log from the master Pi at short intervals to establish up-to-date information on the presence of squirrels at their box. If the arrival of a squirrel was logged, the software started recording a video (top view) from the associated the box, assigned the squirrel as the focal squirrel and started listening for solves. During this time the arrivals and departures of squirrels other than the focal were also recorded. If a solve occurred, the software logged the date, time and side of the lever that was pushed, as well as the focal squirrel’s ID, then operated the two servo motors to reset the lever after a 5s wait. If the focal squirrel departed from the box while additional squirrels were still present, the squirrel that arrived most closely to the focal squirrel became the new focal squirrel. If all squirrels departed the associated antenna, video recording was stopped.

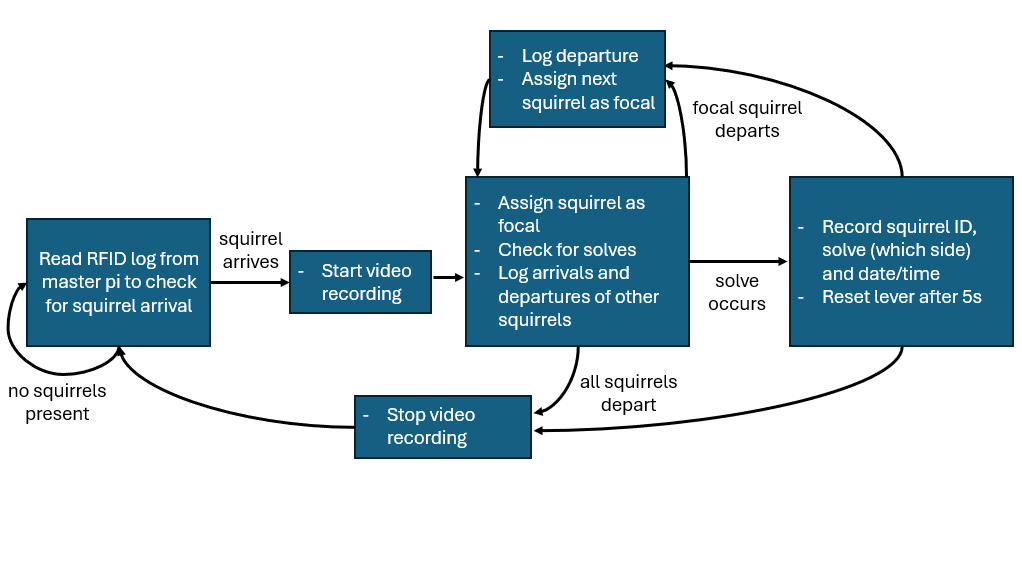

**Figure S1:** Flowchart of puzzle box software

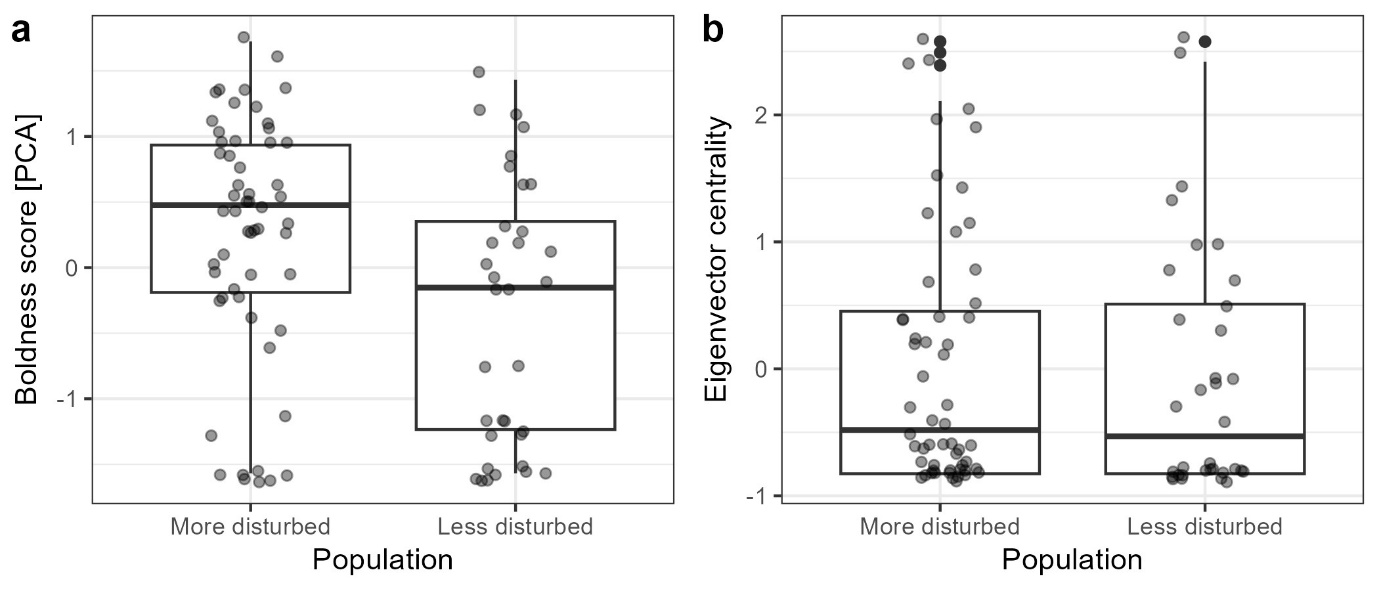

**Figure S2:** Distribution of behavioural types by population for a) boldness; and b) eigenvector centrality as a measure of sociability. Boldness was computed based on scores from a PCA combining trappability and propensity to display fear responses in the trap. Eigenvector centrality was extracted from social networks based on co-feeding events on baited RFID antennae.

**Table S1: Variance Inflation Factors (VIFs) of predictor variables modelling latency to discovery (M1), latency to problem-solving (M2), and task performance (M3)**

|  | M1: Latency to discovery | M2: Latency to solve | M3: Performance |
| --- | --- | --- | --- |
| Mobility | 2.24 | 1.90 | 2.03 |
| Eigenvector centrality | 2.23 | 1.97 | 1.92 |
| Boldness | 1.17 | 1.41 | 1.24 |
| Age | 1.05 | 1.34 | 1.28 |
| Sex | 1.01 | 1.05 | 1.45 |
| Conspecific presence | 1.06 | - | - |
| Population | 1.14 | 1.20 | 1.33 |
| Log # training solves | - | 1.72 | - |
| Observation opportunities (solving) | - | 1.33 | - |

**Table S2: Latency to discovery**

| Predictor | HR | 95% CI (lower) | 95% CI (upper) | z | p |
| --- | --- | --- | --- | --- | --- |
| Mobility (time-varying) | 1.06 | 0.89 | 1.27 | 0.65 | 0.5100 |
| **Boldness** | **1.50** | **1.11** | **2.02** | **2.64** | **0.0083** |
| Eigenvector centrality (time-varying) | 0.99 | 0.84 | 1.17 | -0.09 | 0.9300 |
| Age [J:A] (time-varying) | 0.89 | 0.73 | 1.09 | -1.14 | 0.2500 |
| **Sex [M:F]** | **0.58** | **0.36** | **0.95** | **-2.15** | **0.0320** |
| Conspecific presence (time-varying) | 1.21 | 0.95 | 1.55 | 1.52 | 0.1300 |
| **Population [recreational:trail]** | **2.64** | **1.46** | **4.77** | **3.21** | **0.0013** |

**Table S3: Latency to problem-solving (more disturbed population)**

| Predictor | HR | 95% CI (lower) | 95% CI (upper) | z | p |
| --- | --- | --- | --- | --- | --- |
| Mobility | 1.67 | 0.97 | 2.86 | 1.86 | 0.062 |
| **Boldness** | **2.52** | **1.34** | **4.76** | **2.86** | **0.0042** |
| **Eigenvector centrality** | **2.00** | **1.10** | **3.66** | **2.26** | **0.024** |
| Age [J:A] | 1.06 | 0.33 | 3.42 | 0.10 | 0.92 |
| Sex [M:F] | 0.71 | 0.28 | 1.77 | -0.74 | 0.46 |
| Log # training solves | 1.32 | 0.80 | 2.18 | 1.10 | 0.27 |
| **Obs. opportunities (solving)** | **1.81** | **1.28** | **2.58** | **3.33** | **<0.001** |
| Population [recreational:trail] | 1.04 | 0.39 | 2.79 | 0.08 | 0.94 |

**Table S4: Solving rate**

| Predictor | Estimate | 95% CI (lower) | 95% CI (upper) | z | p |
| --- | --- | --- | --- | --- | --- |
| **Intercept** | **6.46** | **3.33** | **12.93** | **5.59** | **<0.001** |
| **Mobility** | **1.79** | **1.32** | **2.45** | **3.88** | **<0.001** |
| **Boldness** | **1.53** | **1.11** | **2.13** | **2.86** | **0.0043** |
| Eigenvector centrality | 1.11 | 0.81 | 1.53 | 0.69 | 0.49 |
| **Age [J:A]** | **0.55** | **0.30** | **1.01** | **-2.10** | **0.036** |
| **Sex [M:F]** | **2.11** | **1.19** | **3.81** | **2.61** | **0.009** |
| Population [recreational:trail] | 1.27 | 0.69 | 2.25 | 0.82 | 0.41 |

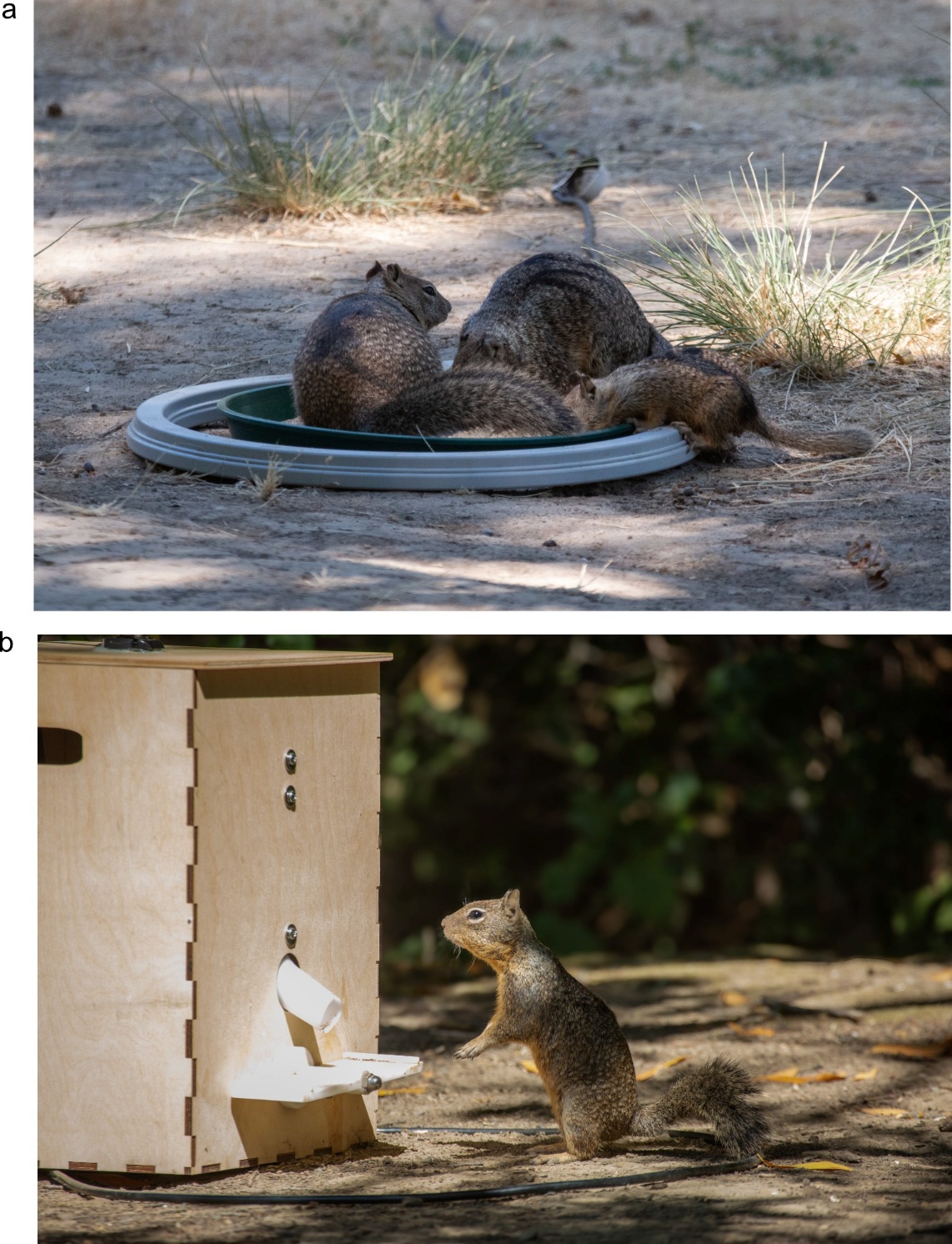

**Figure S3. a:** Three PIT-tagged California ground squirrels forage together on a baited RFID antenna used for mapping social networks based on co-feeding events. **b:** A California ground squirrel approaches an experimental foraging puzzle. These two photos were used to generate illustrations using AI-tools for Figure 2 in the main text.
